# Population genomic analyses reveal a highly differentiated genetic cluster of northern goshawks (*Accipiter gentilis laingi*) in Haida Gwaii

**DOI:** 10.1101/465450

**Authors:** Armando Geraldes, Kenneth K. Askelson, Ellen Nikelski, Frank I. Doyle, William L. Harrower, Kevin Winker, Darren E. Irwin

**Author notes:** Shared first authorship (Armando Geraldes, Kenneth K. Askelson).

## Abstract

Accurate knowledge of geographic ranges and genetic relationships among populations is important when managing a species or population of conservation concern. In the western Canadian province of British Columbia, a subspecies of the northern goshawk (*Accipiter gentilis laingi*) is designated as Threatened under the Canadian Species at Risk Act. Historically, the range of this bird of prey has been ambiguous and its genetic distinctness from the other North American subspecies (*Accipiter gentilis atricapillus*) has not been well established. Given the uncertainty in using morphological traits to assign individual goshawks to these two subspecies, we analyzed genomic relationships in tens of thousands of single nucleotide polymorphisms identified using genotyping-by-sequencing of high-quality genetic samples. This genome-wide analysis revealed a genetically distinct population of northern goshawks on the archipelago of Haida Gwaii and subtle genetic structuring among the remainder of our sampling sites within North America. Following from this analysis, we developed targeted genotyping assays for ten loci that are highly differentiated between the two main genetic clusters, allowing the addition of hundreds of low-quality samples to our analysis. This additional information confirmed that the distinct genetic cluster on Haida Gwaii is restricted to that archipelago. As the *laingi* form was originally described as being based in Haida Gwaii, where the type specimen of that form is from, further study (especially of morphological traits) may indicate a need to restrict this name to the Haida Gwaii genetic cluster. Regardless of taxonomic treatment, our finding of a distinct Haida Gwaii genetic cluster along with the small and historically declining population size of the Haida Gwaii population suggests a high risk of extinction of an ecologically and genetically distinct form of northern goshawk. Outside of Haida Gwaii, sampling regions along the coast of BC and southeast Alaska (often considered regions inhabited by *laingi*) show some subtle differentiation from other North American regions. We anticipate that these results will increase the effectiveness of conservation management of northern goshawks in northwestern North America. More broadly, other conservation-related studies of genetic variation may benefit from the two-step approach we employed that first surveys genomic variation using high-quality samples and then genotypes low-quality samples at particularly informative loci.

## Introduction

Conservation policy and management is becoming increasingly important for the survival of wildlife populations, given the growing impact of human populations on the environment (Robinson, 2006; Barnosky et al., 2011; Côté et al., 2016). An effective tool has been the listing of individual species or subspecies as Endangered or Threatened under the protection of laws such as the Species at Risk Act (in Canada) or the Endangered Species Act (in the USA). This protection has led to the recovery of species such as the peregrine falcon (*Falco peregrinus*; Watts et al., 2015; Ambrose et al., 2016), the Kirtland’s warbler (*Setophaga kirtlandii*; Bocetti et al., 2012), and pink sandverbena (*Ambronia umbellata*; Parks Canada Agency, 2017).

Effective conservation depends on clear identification and classification of true biological entities, enabling identification of the individual organisms and populations that are subject to specific threats, regulations, and management actions. Groups of organisms that look similar to each other but are differentiated genetically, behaviorally, ecologically and/or physiologically (Bickford et al., 2007; Toews & Irwin, 2008; Pulido-Santacruz et al., 2018) pose a particular conservation challenge for policy-makers. Such “cryptic” biodiversity is important to identify and manage properly, as the extinction of one of the cryptic forms would result in the loss of important biological variation. Likewise, when the boundaries between forms that are only subtly differentiated (e.g., two subspecies or otherwise differentiated populations) are unclear, efforts to conserve one of those forms are greatly complicated.

Northern goshawks (*Accipiter gentilis*) living along the coast of British Columbia, Canada, and southeast Alaska, USA, provide an example of a species with unclear boundaries between forms that differ in their conservation status listing. Taverner (1940) determined that northern goshawks from the Haida Gwaii archipelago (called the Queen Charlotte Islands at the time) were more darkly colored than those from the mainland, whereas northern goshawks from Vancouver Island were “more variable and less plainly characterized.” Taverner (1940) formally designated this darker form with the subspecies name “*A. g. laingi*” (also, the “Queen Charlotte Goshawk”), using a specimen from Haida Gwaii (from Masset, on Graham Island) as the type specimen. The other major North American subspecies of northern goshawk is *A. g. atricapillus*, which ranges across much of the continent. Since Taverner’s (1940) work, the exact ranges of these two subspecies along the coast of BC, southeast Alaska, and the state of Washington has been under debate. About one-third of northern goshawks on Vancouver Island and within the Alexander Archipelago of southeast Alaska have been described as having a dark appearance approaching (but not completely matching) the darkness of those on Haida Gwaii (COSEWIC, 2013; Titus et al., 1994; Webster, 1988), providing equivocal evidence as to whether the range of *laingi* should be considered to include those areas. Given this ambiguity from the appearance of the northern goshawks, biologists have turned to habitat modeling (based in part on radio-telemetry data) to more specifically delineate the inferred range of *laingi*. For example, the Northern Goshawk *Accipiter gentilis laingi* Recovery Team (2008) mapped the range of *laingi* as corresponding to “the distribution of wet Coastal Western Hemlock (CWH) biogeoclimatic subzones/variants and the Coastal Douglas-fir (CDF) biogeoclimatic zone.” This area includes much of coastal British Columbia, southeast Alaska, and western Washington. The small and declining population of northern goshawks within this region (estimated at roughly 1200 individuals within British Columbia; COSEWIC, 2013), as well as threats such as habitat loss, have led to *laingi* being listed as Threatened under the Canadian Species at Risk Act (COSEWIC, 2000, 2013). However, continued debate over which individual northern goshawks should be considered *laingi* versus *atricapillus*, and hence which geographic areas should be considered within the range of each subspecies, has complicated conservation policy and management (COSEWIC, 2013).

Studies of molecular genetic variation can be of great use in revealing natural biological groups and in assigning individuals to those groups (Wagner et al., 2013; Mason & Taylor 2015). Previous genetic research in northern goshawks was inconclusive in terms of clarifying the range of *laingi*. Sonsthagen et al. (2012) concluded that patterns in both mitochondrial DNA and microsatellites were consistent with gene flow among sites along the BC coast and southeast Alaska, but the study did not include northern goshawks from outside of this region. Most analyses of genetic variation in a conservation context have used a small set of molecular markers (e.g., mitochondrial DNA, which is inherited as a single unit from the mother; or a small number, e.g. 5-20, of microsatellite loci). These approaches may not reveal true population differences because the portion of the genome that differs between biological groups can be quite limited, even between groups that are morphologically distinct (Mason & Taylor 2015; Toews et al., 2016b). Recent advances in genomic sequencing allow for a much more comprehensive survey of genetic variation across the genome, such that small portions of the genome that differ between groups can be detected (Toews et al., 2016b).

Here we used a two-stage analysis that first surveys genome-wide variation among high-quality genetic samples and then examines genotypes of highly informative loci in a much larger set of low-quality genetic samples. Firstly, we used genotyping-by-sequencing (GBS; Elshire *et al*., 2011) to survey geographic variation in northern goshawks at tens of thousands of variable sites (i.e., single nucleotide polymorphisms [SNPs]) across their genome. Our sampling includes goshawks from Haida Gwaii, Vancouver Island, coastal and interior British Columbia, southeast and northern Alaska, Washington state, the midwestern and eastern USA, and Europe. To assess congruence between nuclear and mitochondrial DNA patterns, we analyzed variation in mitochondrial DNA control region sequences. Secondly, given that many of the northern goshawk samples are from shed feathers or museum skin samples that provided DNA of insufficient quality for the GBS analysis, we developed assays for 10 markers with high frequency difference between the two main genetic clusters revealed by our genome-wide analysis. These assays were then used to genotype many low-quality DNA samples. We propose that this general two-step approach could be used to better clarify the ranges of other groups of conservation concern and to determine the genetic ancestry of otherwise difficult-to-identify individuals.

## Materials and Methods

### 1. Sampling and DNA extraction

A total of 433 northern goshawk (*Accipiter gentilis*) samples from across North America were used in this study. In addition, nine samples of the European subspecies of northern goshawk (*A. g. gentilis*) and five samples from other members of Accipitridae (two Cooper’s hawks [*A. cooperii*], one sharp-shinned hawk [*A. striatus*], one red-tailed hawk *[Buteo jamaicensis*] and one bald eagle [*Haliaeetus leucocephalus*]) were also included in this study (Supporting Table 1). The tissue type used for DNA varied; we extracted shed feathers (usually collected from nests or from the ground near nests), blood (collected from live birds), muscle tissue (from dead birds, now museum specimens), and toe pads (from museum study skins). All samples used in the data analysis were from different individuals (see below for details).

DNA was extracted using a standard phenol-chloroform protocol following overnight digestion at 55°C with 0.3 μg of proteinase K in 400 μL of lysis buffer (0.1 M Tris, 5 mM EDTA, 0.2% SDS, 0.2 M NaCl, pH 8.5). In samples consisting of shed feathers, a separate digestion protocol was used (following Bayard de Volo et al., 2008) before DNA extraction. DNA was resuspended in 50-100 μL of 1X TE buffer and quantified in a Qubit fluorometer (Invitrogen) following the manufacturer’s instructions. The quality of DNA yielded from the different types of source material varied considerably, with blood and tissue producing the highest quality DNA in the largest quantities. Feathers and toe pad samples produced mostly degraded DNA (i.e., broken into relatively small fragments, hence appearing as a low-molecular-weight smear after electrophoresis), but toe pads typically produced more DNA than feathers did. Because the majority of our samples were from shed feathers, and because they produced mostly degraded DNA, we used a two-step approach in terms of methodology: we conducted GBS on our high-quality tissue samples and Taqman genotyping assays of informative loci on a broader set of samples of varying DNA quality (described below). Some feather samples from critical areas like Haida Gwaii and Vancouver Island were included on our GBS plates but in general they performed poorly (described below).

### 2. DNA sequencing

#### a) Genotyping-by Sequencing (GBS)

We used a reduced representation genome sequencing method, GBS (Elshire et al., 2011), following modifications as in Alcaide et al. (2014) and specified in detail in Toews et al. (2016a). We made the following minor modifications to the protocol in Toews et al. (2016a): a) we used 10 μL of cleaned DNA fragments from the ligation reaction for a PCR of 18 cycles; b) after Qubit quantification of PCR products, 100 ng of each PCR product was added to the pool which was then concentrated in a SpeedVac (Savant DNA110, TheromoFisher) and run on 5 lanes of a 2% agarose gel; and c) we selected fragments within a size range of 400-500 bp. Two GBS libraries, each containing up to 96 individually barcoded samples, were prepared and submitted for paired-end sequencing on an Illumina HiSeq 2500 (at McGill University and Génome Québec Innovation Centre). For inclusion in GBS we prioritized samples for which extractions yielded high molecular weight DNA, but we also included eleven samples whose extractions yielded considerably degraded DNA because they represented birds from important areas for this study (Haida Gwaii, *n* = 7; Vancouver Island, *n* = 1; mainland British Columbia, *n* = 2; and coastal mainland British Columbia, *n* = 1). The starting materials for DNA extraction for these degraded samples were shed feathers collected near nests (*n* = 10) or toe pads (*n* = 1). These 11 samples were submitted for sequencing multiple times (with different barcodes) to try to obtain enough sequence data for our analyses. Across the two libraries, a total of 172 barcodes were used for 160 samples (plus 4 blanks, two no-DNA blanks and two no-barcode blanks, and 16 samples from other projects; Supporting Table 1). All resulting DNA sequences will be deposited in the NCBI SRA under accession number #### (following peer-reviewed journal publication).

#### b) mtDNA sequencing

We used the primers L16064 and H15426 (Sonsthagen et al., 2004) to PCR amplify and Sanger sequence 578 bp of the mitochondrial control region (between positions 1169 and 1747 of the complete mitochondrial sequence of *A. gentilis* [GenBank: AP010797.1]) in 125 *A. gentilis*, two *A. cooperii*, one *A. striatus*, one *Buteo jamaicensis* and one *Haliaeetus leucocephalus* (Supporting Table 1). PCR amplifications were carried out in a total volume of 25 μl, with 0.5 μM of each primer, 0.2 mM of dNTP’s, 2 mM of MgCl_2_, 0.5 units of recombinant *Taq* DNA polymerase (Invitrogen), and between 2 and 20 ng of DNA. Cycling conditions were: 94° for 3 minutes; 35 cycles of 94° for 45 seconds, 51° for 30 seconds, and 72° for 1 minute; and a final extension step at 72° for 10 minutes. PCR products were sent out for purification and Sanger sequencing at Macrogen (USA) using primer H15426. Mitochondrial DNA (mtDNA) sequences were visually inspected and manually aligned with BioEdit (Hall, 1999). The bald eagle and red-tailed hawk sequences were so divergent that they could not easily be aligned, and they were not considered further. The final sequence alignment will be deposited in NCBI under accession number #### (following peer-reviewed journal publication).

### 3. GBS Sequencing, filtering, and analysis

GBS sequencing of the two libraries resulted in over 1.12 billion reads in total. To analyze these reads we followed the scripts available at https://doi.org/10.5061/dryad.t951d from Irwin et al. (2016) for GBS read processing and mapping. In brief, we demultiplexed the raw GBS reads from the two barcoded libraries using a custom perl script from Baute et al. (2016) that separated reads according to the barcodes for each sample (Supporting Table 1), removed barcode and adaptor sequences and removed sequences shorter than 30 bp. A fraction of reads in each sequencing lane could not be assigned to our barcodes and were discarded (13.8% and 15.3% of reads in each sequencing lane). We then trimmed the reads with Trimmomatic-0.36 (Bolger et al., 2014) with options TRAILING:3, SLIDINGWINDOW:4:10, MINLEN:30. We aligned the trimmed reads for each barcoded sample to the bald eagle reference genome (Warren et al., 2014, assembly GCF_000737465.1) using BWA-MEM (Li & Durbin, 2009) with default settings. This reference genome is from a male bird (hence, no W chromosome sequence is represented), and it is assembled into 1,023 scaffolds which have not been anchored to chromosomes. An average of 5.59 M reads per sample (range 92 reads to 26.33 M reads) were used for mapping to the bald eagle reference genome with BWA-MEM, and an average of 4.89 M reads per sample (range 55 reads to 25.49 M reads) were successfully mapped. Despite mapping reads to a distantly related species (bald eagle to northern goshawk estimated divergence time is 31 MYA http://www.timetree.org/), across all samples an average of 84.8% of reads were successfully mapped, and for 134 samples, more than 95% of reads were successfully mapped. As expected, the sample with the highest fraction of reads mapped was from our bald eagle (98.7% reads mapped). The largest source of variation in the number of reads generated and number of reads mapped per sample appeared to be the source material used for DNA extraction. The two most common sources of DNA in our GBS analysis were muscle tissue preserved in ethanol (112 samples) and shed feathers (feathers found near nests; 33 samples). While on average both types of material generated large numbers of reads (4.61 M for feathers and 5.92 M for tissue), the average number of reads mapped was much lower for feathers (1.84 M) than for tissue (5.75 M). Three samples (Supporting Table 1) had less than 100 K reads mapped and were dropped from further analyses. Remaining analyses were performed only on the remaining 157 samples.

Despite most reads having been mapped to the bald eagle genome, our requirement that reads map with high quality (MAPQ≥20) meant that on average only 56.5% of mapped reads from these 157 samples were used for further analysis. The evolutionary distance between northern goshawks and the bald eagle leads to lower mapping quality for northern goshawk GBS reads (e.g., for sample NGAK020, only 55.6% reads mapped with MAPQ ≥ 20; Supporting Figure 1) in comparison with our bald eagle GBS reads (84.6% of bald eagle reads mapped with MAPQ ≥ 20).

We used Picard (http://broadinstitute.github.io/picard/) and SAMtools (Li et al., 2009) to generate for each individual a BAM file containing its sequence information. Reads from samples run with multiple barcodes were merged into a single file at this stage. We called genotypes for one sample at a time using GATK v3 (McKenna et al., 2010) with the function HaplotypeCaller and then generated a single VCF file with the GATK function GenotypeGVCFs with all 157 samples for which more than 100 K reads had been mapped with high confidence (Supporting Table 1). Unlike Irwin et al. (2016), we a) did not realign around indels with functions RealignerTargetCreator and IndelRealigner as this is discouraged with recent versions of GATK, b) changed option “-max_alternate_alleles 2” to “–max_alternate_alleles 4” in HaplotypeCaller, c) did not use the options “–allSites” and “-L” in GenotypeGVCFs, and d) used option “-hets 0.01” in GenotypeGVCFs.

The resulting VCF file was further filtered using VCFtools v0.1.11 (Danecek et al., 2011) to remove insertion and deletion polymorphisms and loci that were not biallelic. We used a custom perl script (Owens et al., 2016) to filter out loci with observed heterozygosity of 0.6 or greater as these are likely the result of paralogous variation instead of allelic variation. At this stage, we used VCFtools to remove from further analysis 22 samples that had more than 80% missing loci in the dataset prior to heterozygosity filtering. Shed feathers were 22 of the 25 (88.0%) eliminated samples due to low reads or high missing data, and only comprised 12 of the 135 (8.9%) remaining samples (Supporting Table 1). Additionally, 554 loci suspected of being sex linked (Supporting Table 2 and Supplemental File 1) were removed from analysis, and seven samples identified as having close kin relationships with other samples in the study were also excluded from further analysis (Supporting Tables 3 and 4 and Supplemental File 1).

The resulting dataset included 128 individuals (Table 1 and Figure 1) and 2,885,805 SNPs. For specific analyses of population structure, this dataset was filtered in various ways (described below with each analysis); all involved filtering (using VCFtools) to include only SNPs with less than 30% missing genotypes and genotype quality of 10 or higher. We filtered out SNPs with rare alleles by either eliminating SNPs with rare alleles that occurred only once (i.e., eliminating singletons) or by keeping only SNPs with a minor allele frequency of 0.05 or higher. Different analyses included either all individuals in the study (*n* = 128), only goshawks (*n* = 124), only North American goshawks (*n* = 119), or only North American goshawks excluding those from Haida Gwaii (*n* = 107). In some analyses (see below), we filtered to include only unlinked SNPs, using the R package SNPrelate (Zheng et al., 2012).

**TABLE 1.** Numbers of samples per population for which data was used in the GBS, mtDNA and Taqman data analyses. Sample details in Supporting Table 1.

| Population | Population code | GBS | mtDNA | TaqMan |
| --- | --- | --- | --- | --- |
| Alaska - North | AK | 26 | 27 | 43 |
| Alaska - Alexander Archipelago | AA | 17 | 18 | 20 |
| Arizona | AZ | 0 | 0 | 9 |
| British Columbia - Interior | BC | 11 | 16 | 118 |
| British Columbia - Coastal | BCc | 5 | 5 | 45 |
| British Columbia - Haida Gwaii | HG | 12 | 7 | 14 |
| British Columbia - Vancouver Island | VI | 6 | 7 | 79 |
| California | CA | 0 | 0 | 4 |
| Eastern United States | East | 29 | 28 | 40 |
| Washington state | WA | 13 | 13 | 14 |
| Europe | Eur | 5 | 4 | 0 |
| Cooper's hawk | OUT-CH | 1 | 2 | 0 |
| Sharp-shinned hawk | OUT-SS | 1 | 0 | 0 |
| Bald eagle | OUT-BE | 1 | 0 | 0 |
| Red-tailed hawk | OUT-RT | 1 | 0 | 0 |
| Total |  | 128 | 127 | 386 |

**FIGURE 1.**
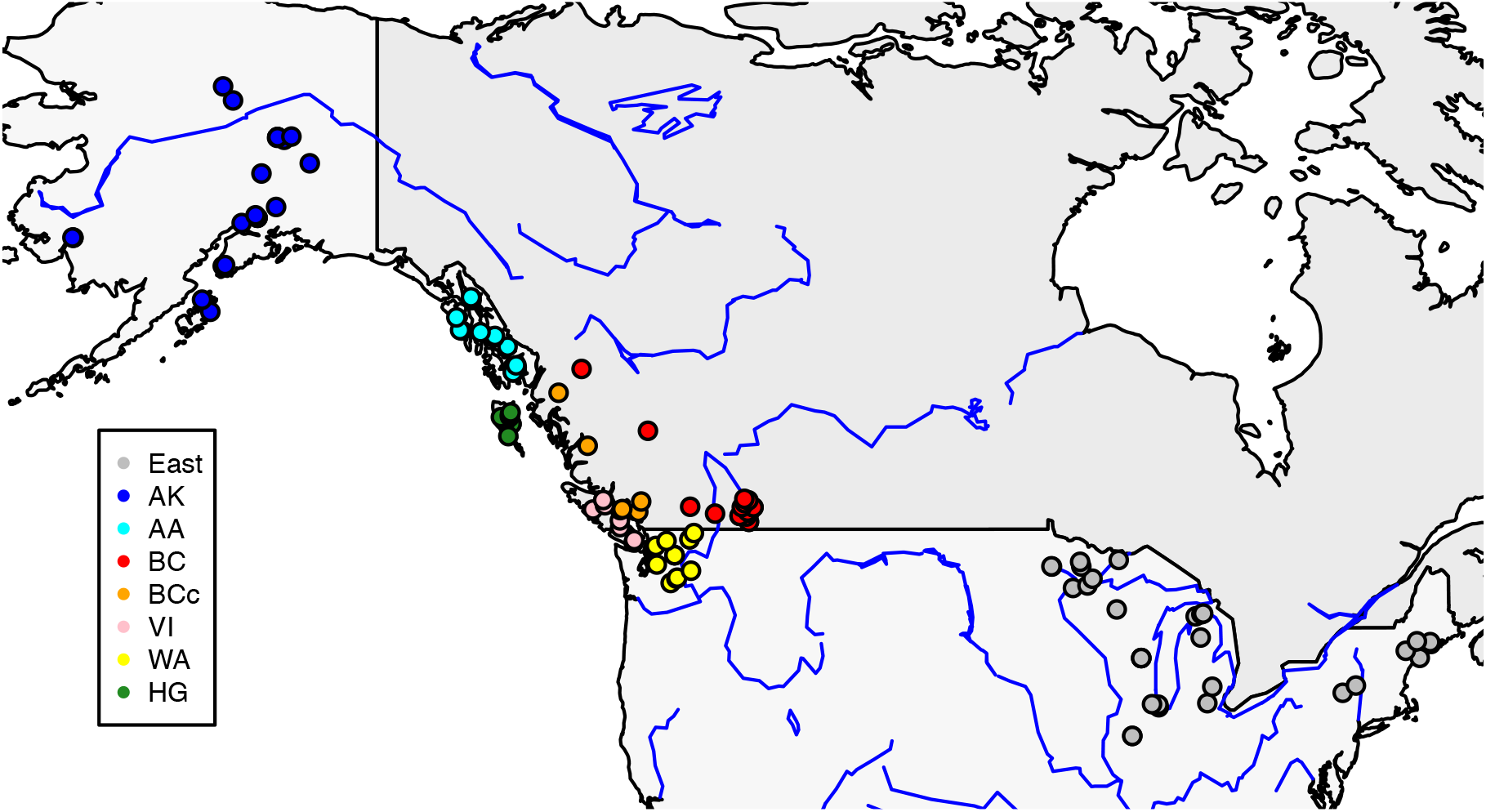
Map of North America showing the provenance of northern goshawk samples on which GBS and/or mtDNA analyses were performed. Sample sizes can be found in Table 1. Population abbreviations are: East (Eastern United States), AK (Alaska north), AA (Alexander Archipelago of southeast Alaska), BC (interior mainland British Columbia), BCc (coastal mainland British Columbia), VI (Vancouver Island), WA (Washington State) and HG (Haida Gwaii).

### 4. Population Structure and Genetic Differentiation

The overall relationships between all individuals in the genus *Accipiter* were estimated with an unrooted phylogenetic network with uncorrected p-distances (Nei and Kumar 2000) using SplitsTree4 V4.14.5 (Huson & Bryant, 2006) after eliminating singletons and the two non-*Accipiter* spp. individuals in VCFtools (*n* = 125,818 SNPs and 126 samples). For the mtDNA dataset, we estimated a Neighbour-Joining tree (Saitou & Nei, 1987) using the alignment of all *Accipiter* spp. sequences and uncorrected p-distances (Nei & Kumar, 2000) in MEGA X (Kumar et al., 2018). We also produced a haplotype network with the median joining algorithm (Bandelt et al., 1999) in the program Network v5.0 (http://www.fluxus-technology.com) for the North American northern goshawk sequences.

We used two complementary approaches to inferring patterns of population structure among North American goshawks using the GBS dataset, after filtering to unlinked SNPs with minor allele frequency (MAF) > 0.05 (*n* = 6,058 SNPs). First, we performed principal components analyses (PCA) with the R package pcaMethods (Stacklies et al., 2007), with missing genotypes imputed with svdImpute. Second, we used Admixture v1.3.0 (Alexander et al., 2009) to estimate ancestry proportions for each sample. Admixture is a clustering program that, like Structure (Pritchard et al., 2000), models the probability of the observed genotypes using ancestry proportions and population allele frequencies. Instead of a Bayesian approach, however, Admixture uses a Maximum Likelihood approach resulting in much faster runs. We ran five replicates of Admixture allowing for the number of clusters (*K*) in the model to vary from 1 to 9, and we terminated each run when the difference in log-likelihood between successive iterations fell below 1×10^−9^. We chose the *K* that minimized cross-validation error and hence best fit the data (Alexander et al., 2009).

To test for admixture between all northern goshawk sampling regions (*n* = 124 samples), we calculated the f3 statistic (Reich et al., 2009) using Treemix v1.3 (Pickrell & Pritchard 2012) on our SNP dataset with singletons excluded and no initial filtering based on linkage (*n* = 85,039 SNPs). The f3 statistic can be used to test whether a population, e.g., *X*, is the result of admixture between two others, e.g., *Y* and *Z*. We calculated this f3 statistic for all 336 possible combinations of three sampling regions (choosing from the 8 sampling regions) in the dataset. A significantly negative value for the f3 statistic (*X*; *Y*, *Z*), indicates that sampling region *X* is admixed between *Y* and *Z*, whereas significantly positive values indicate no admixture. To account for linkage between nearby SNPs we used the command “-k 50” in the estimation of f3 statistics so that SNPs were blocked in windows of 50 consecutive SNPs ordered assuming synteny with the bald eagle.

Given that our results (see below) indicated population differentiation between Haida Gwaii and elsewhere, and to a lesser degree between coastal regions and elsewhere, we used assignment tests to explore the degree to which our genomic dataset can be used to predict geographic origin of a sample. This was done by applying Discriminant Analysis of Principal Components using the R package adegenet (Jombart, 2008) and using cross-validation values as probabilities of correct assignment. We trained our data sets with 60% of the data and tested them with the remaining 40%. Tests of assignment probability were performed across 10 PCA axes and for each PCA axis tests were replicated 20 times. Missing genotypes were imputed with the mean allele frequency for each SNP. We averaged correct assignment proportions across the first 10 PCA axes and across the 20 replicates.

### 5. Inbreeding

To test whether there is more inbreeding in some regions compared to others, we calculated the average genome-wide inbreeding coefficient (*F*) for each individual within each sampling region using the method of moments implemented in VCFtools (option “–het”, Danecek et al., 2011). We did this for each North American sampling region separately after first filtering out singletons (resulting in 24,322 SNPs).

### 6. Divergence between species/subspecies

To quantify patterns of relative population differentiation, we used VCFtools (command “–weir-fst-pop”, after first eliminating singleton SNPs) to estimate overall pairwise weighted *F*_ST_ between sampling regions following Weir & Cockerham (1984).

For the same groups we also estimated net nucleotide divergence, *D*_A_ (Nei, 1987). *D*_A_ is defined as *D*_XY_ - 0.5(*D*_X_ + *D*_Y_), where *D*_XY_ is the average pairwise nucleotide distance between groups and *D*_X_ and *D*_Y_ are the average pairwise nucleotide distances within groups. *D*_XY_, *D*_X_, and *D*_Y_ were estimated across the 40 largest scaffolds, representing 592,096,274 bp, or 50.24% of the published genome sequence of the bald eagle, using the GATK pipeline and a custom R script described in Irwin et al. (2016). We retained invariant SNPs (using the “-allSites”option in GATK) and included SNPs with up to 60% missing data. For each statistic (*D*_XY_, *D*_X_, and *D*_Y_) we generated estimates in windows (each containing 5000 sequenced bp) across the 40 scaffolds, and we then averaged the windows together for a single overall estimate between each pair of populations. For the mtDNA dataset sequence-based *F*_ST_ (Hudson et al., 1992) was estimated in DNAsp v6 (Rozas et al., 2017) and *D*_A_ in MEGA X (Kumar et al., 2018). To account for recurrent mutations, we used a Maximum Composite Likelihood model (Tamura et al., 2004) with rate variation among sites modeled with a gamma distribution (shape parameter = 0.05) and taking into account composition bias among sequences (Tamura & Kumar 2002).

We used the differentiation estimates between Haida Gwaii and the remaining North American sampling regions to get a rough estimate of the time since they started diverging using the expectation from the neutral theory that *D* = 2*μt*, where *μ* is mutation rate, *t* is time, and *D* is genetic distance. We used *D*_A_ as a proxy for net genetic distance *D*, an approach that takes into account within-group variation. We used the nuclear DNA substitution rate as estimated from *Ficedula* flycatchers (*μ* = 2.30×10^−9^ per year per base pair; Smeds et al., 2016), and we estimated the mitochondrial DNA mutation rate as *μ* = 2.9×10^−7^ per year per base pair assuming an estimated divergence time between northern goshawks and Cooper’s hawk of 11.1 MY (http://www.timetree.org/).

### 7. Ancestry informative assays

Our GBS results showed two main genetic clusters of North American goshawks (see Results below): one consisting of individuals from Haida Gwaii and one consisting of individuals from all other North American sampling sites. To determine to which of these clusters the remaining samples in our study belong, we developed SNP genotyping assays for a subset of SNPs that are highly differentiated between the two clusters. Using VCFtools (Danecek et al., 2011), we estimated *F*_ST_ for the 9,850 SNPs (no LD pruning) between samples from Haida Gwaii (*n* = 12) and the remaining North American northern goshawk samples (*n* = 107). Loci with MAF < 0.05 were eliminated because these could not have large allelic frequency differences between populations. We then inspected each SNP in descending order of their *F*_ST_ rank to determine their suitability for designing custom TaqMan (Applied Biosystems) SNP genotyping assays. We selected 11 SNPs among those with highest *F*_ST_ for which we had at least 30 base pairs of sequence on either side of the target SNP, for which there were no other variants in the flanking sequence (or if present in the entire dataset, they had to be rare), and that were all from different contigs in the bald eagle reference genome (Supporting Table 5).

For each locus, samples were genotyped in 384-well plates following the manufacturer’s instructions: 2.5 μl TaqMan genotyping master mix 2X, 0.25 μl TaqMan assay 20X, 1 μl DNA (concentration between 1 and 5 ng/μl) and 1.25 μl water. Genotyping was performed in a Viia7 Real-Time PCR system (Applied Biosystems) with the following conditions: 95° for 10 minutes, and 40 cycles of: 95° for 15 seconds and 60° for one minute. We called the genotypes for each sample at each locus by visual inspection of the plots of the ΔRn values of each allele. The Rn value is the reporter dye (FAM) signal normalized by the fluorescence signal of the ROX dye, and ΔRn is Rn minus the baseline.

We performed a trial genotyping assay for each of the 11 loci with a small subset of samples for which we had genotypes at these loci from the GBS data. Genotyping was successful at 10 of the 11 TaqMan loci. Each of the 10 successful loci were then genotyped in the entire sample set in two 384 plates each containing 32 reference samples (Supporting Table 1). These 32 samples were selected from our GBS samples so that for each locus there were multiple observations of each genotype (homozygous allele 1, heterozygous, and homozygous allele 2).

These 10 assays were used to genotype 444 samples from our entire sample collection, which included large numbers of shed feathers and toe pads from museum skins for which we were only able to extract small amounts of highly degraded DNA. These data will be deposited in Dryad accession number #### (following peer-reviewed journal publication). When multiple feathers were available from a single nesting territory only one was used for the GBS analysis, but for the TaqMan assays multiple feathers were used for 13 territories. In only one case, different feathers from the same territory were kept for further analyses because their multilocus genotypes differed. We successfully genotyped 386 North American northern goshawks for 7 or more of the 10 loci (Supporting Table 6). The genotyping rate was high (range 89.6 to 99.7%), and the discrepancy rate between the GBS and TaqMan datasets was low (0.79%; i.e., we observed eight genotype discrepancies between datasets out of a total of 1,013 genotypes). Seven discrepancies consisted of a heterozygote genotype for TaqMan and a homozygote genotype for GBS. One discrepancy consisted of homozygote genotypes for different alleles (Supporting Table 6 and Supporting Figure 2).

### 8. Distributions of main genetic clusters

We used the R package HIest (Fitzpatrick, 2012) to estimate in a maximum likelihood framework the ancestry index (*S*) and interclass heterozygosity (*H*) of each North American goshawk sample for which there were at least 7 TaqMan genotypes. HIest allows for the estimation of *S* and *H* even in the absence of diagnostic markers, as is the case here, given that for each locus we have allele frequency estimates from reference populations. For reference populations we selected from our GBS dataset the 12 individuals from Haida Gwaii and the 28 individuals from the Eastern United States for which we also had TaqMan data. In order to compare estimates of ancestry proportions from our TaqMan data to our GBS dataset we used Admixture with the genotyping results from these 10 loci.

## Results

### Overall species relationships

Estimated phylogenetic relationships among the *Accipiter* samples used in this study are illustrated in Figure 2, based on both the GBS dataset and the mtDNA dataset. These networks both depict, as expected, sharp-shinned and Cooper’s hawks (i.e., the other two *Accipiter* species) as outgroups to northern goshawks. Within northern goshawks, both nuclear and mitochondrial datasets are congruent in showing a pattern of close relationships within North America compared to the distant relationship between North American and European populations. Analyses of the nuclear dataset with Admixture and PCA reveals a similar pattern of strong differentiation between European (*A. g. gentilis*) and North American samples of northern goshawks (Supporting Figure 3).

**FIGURE 2.**
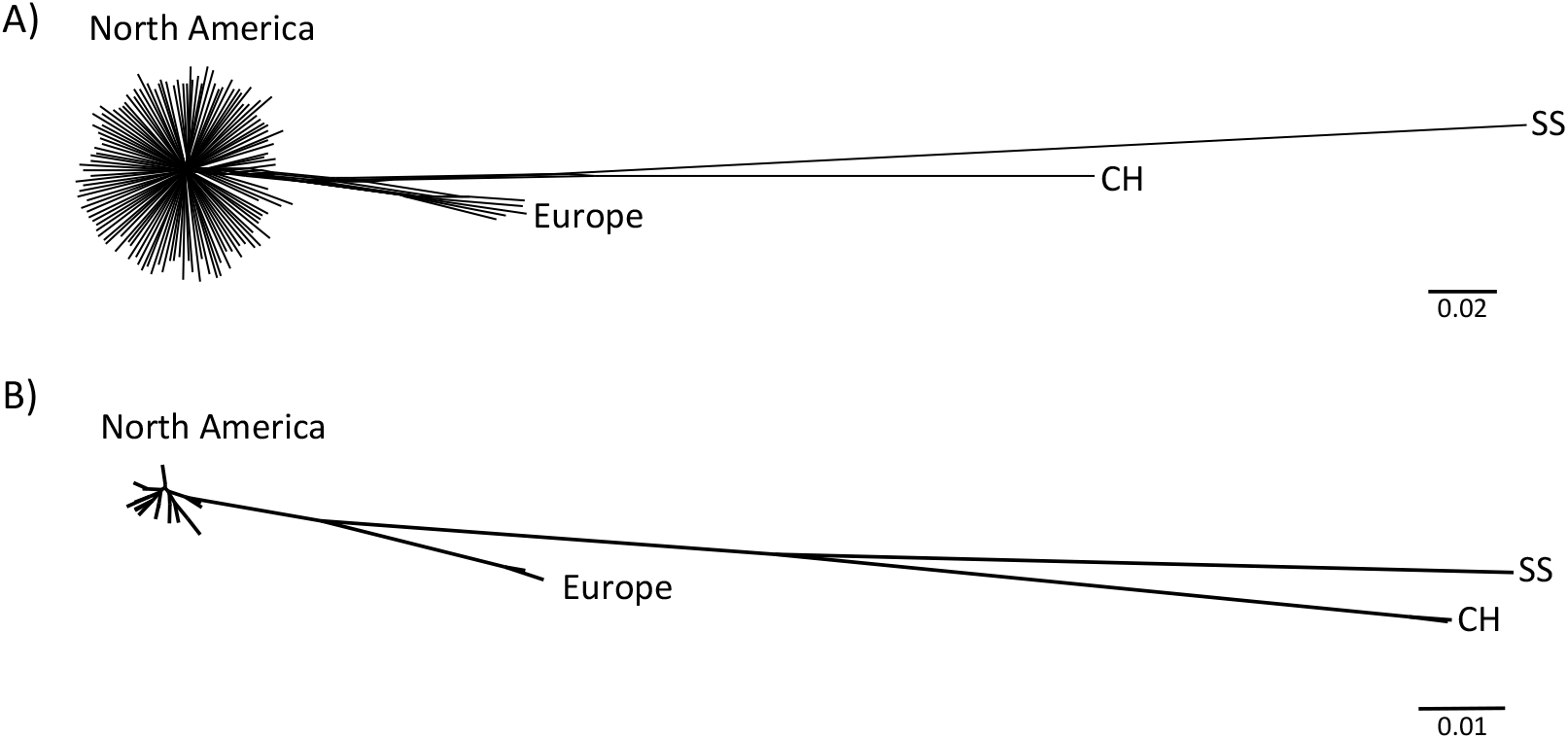
Phylogenetic relationships among *Accipiter* spp. sampled in this study. A) Unrooted network for the nuclear GBS dataset including 126 samples and 125,818 SNPs (singletons excluded) and B) Neighbour-Joining network for the mtDNA dataset including 128 samples. Both were estimated with uncorrected *p*-distances. SS: sharp-shinned hawk; CH: Cooper’s hawk. Sample details in Table 1 and Supporting Table 1.

### Population structure within North American northern goshawks

Analysis of the North American northern goshawk samples clearly reveals two genetic clusters: one that includes all individuals from Haida Gwaii (HG), and a second containing all remaining North American individuals (see population abbreviations and sample sizes in Table 1 and Figure 1). This pattern is illustrated using both a Principal Components Analysis (PCA) and an Admixture analysis (Figure 3 and Supporting Table 7). The first PCA axis, explaining 5.8% of the genotypic variation, separates HG from all other sampling regions (Figure 3). Similarly, in the Admixture analysis, where *K* = 2 was the *K* value with the greatest support (Supporting Figure 4), 11 out of 12 HG individuals have an estimated 100% ancestry in one ancestral population (with the remaining HG individual having 89% ancestry in that same population). All individuals from other sampling regions are estimated to have a majority of their ancestry (ranging from 81% to 100%, mean of 96%) from the second ancestral population. Only 17 individuals (or 16%) are estimated to have 10% or more ancestry in the HG cluster: 15 from the Alexander Archipelago (AA) of southeast Alaska, one from coastal British Columbia (BCc), and one from Vancouver Island (VI). These 17 individuals also have the lowest PC1 values of any samples outside of HG (Fig. 3A).

**FIGURE 3.**
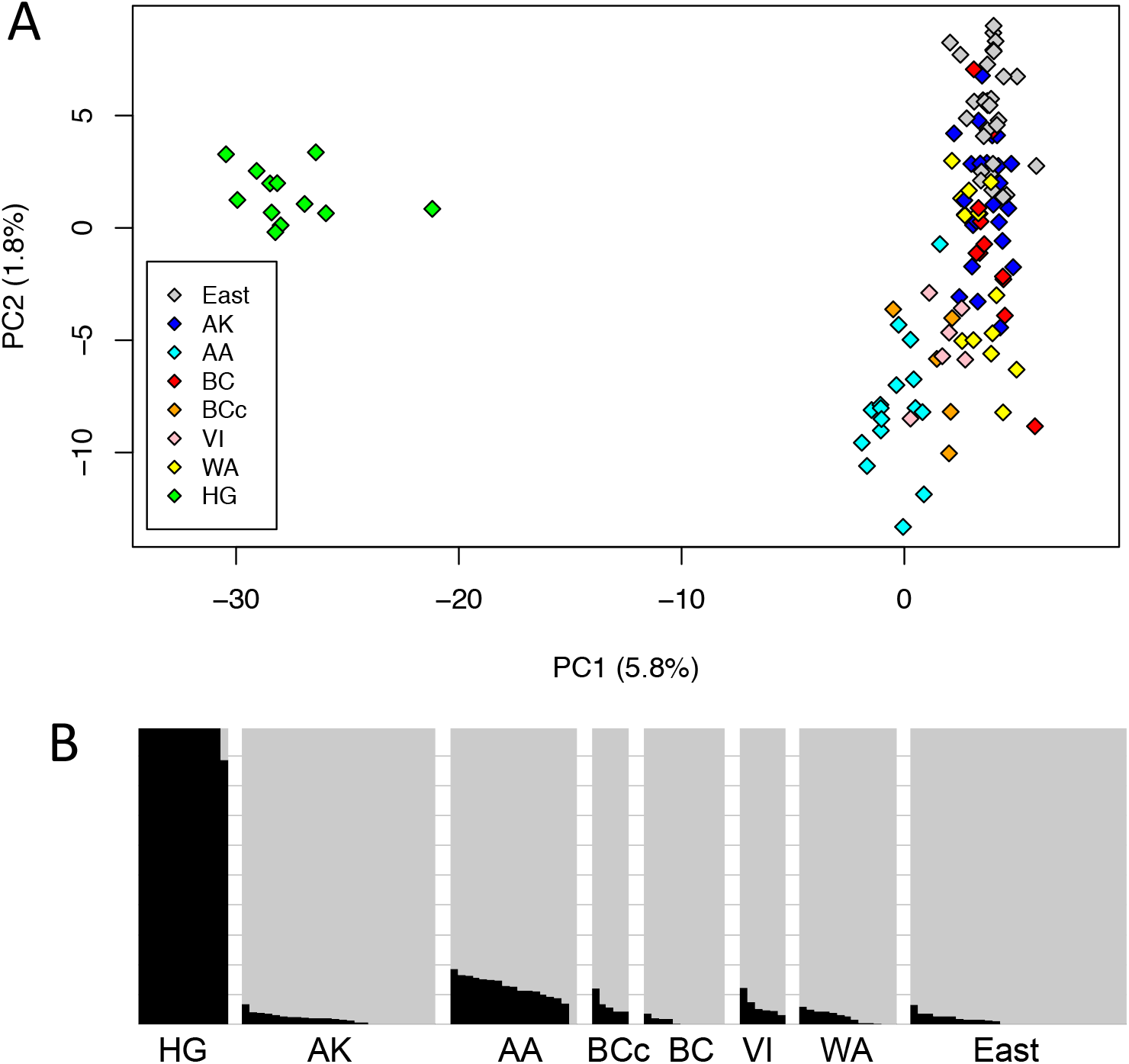
Population structure of the northern goshawk samples from North America based on genomic relationships estimated using GBS. Principal Components Analysis (A) and Admixture analysis (B) were performed on 6,058 unlinked SNPs with minor allele frequency of 0.05 or above. Population names and sample sizes can be found in Table 1. In panel B, each vertical column represents a different sample. The height of black and gray indicates the estimated proportions of a sample’s genome derived from two inferred ancestral populations. Further PC axes did not reveal any obvious geographic trends and *K* = 2 was the value that minimized the cross-validation error in Admixture.

The second axis of variation in the PCA explains only 1.8% of the genotypic variation in the data, and unlike what is seen in PC1, no sampling regions cluster together to the exclusion of any others, and no large gaps in the distribution of samples are observed. Despite this, some subtle differences in average position of sampling regions along PC2 can be detected. Samples from AA (*n* = 17), VI (*n* = 6), and BCc (*n* = 5) tend to have negative values along PC2, whereas all samples from the Eastern USA (East, *n* = 29) have positive values along PC2. Interior BC (BC, *n* = 11), Alaska north (AK, *n* = 26), and Washington state (WA, *n* = 13) tend to have intermediate values of PC2. Hence there is some geographic structure within the non-HG cluster, but it is small compared to the very clear differentiation between HG and elsewhere. Further evidence for only weak North American population structure outside of HG comes from the fact that *K* = 2 (as above) is the most-supported *K* in the Admixture analyses. Plotting the Admixture results for *K* = 3 (Supporting Figure 5) again reveals HG as one population and most remaining samples are represented as mixtures between two additional populations with no apparent geographic pattern. The overall PCA pattern is similar when rare alleles are considered (i.e., including all unlinked SNPs that are observed at least twice; Supporting Figure 6), but the Admixture analysis detects no population structure (i.e., one is the *K* value that minimizes the cross-validation error). We repeated these analyses with HG excluded, and again results suggest some weak population structuring in North America outside of HG (Supporting Figure 7 and Supporting Figure 8).

These general patterns are further supported by cross-validation tests using discriminant analysis of principal components (DAPC). When testing assignment to HG vs. elsewhere in North America, cross-validation correctly assigns individuals 96.2% of the time. Without HG in the dataset, a test of assignment to coastal (that is, AA, VI, and BCc treated as a single group) vs. elsewhere in North America correctly assigns individuals 83.3% of the time (note that random assignment to two groups would be right 50% of the time). This suggests some differentiation of these regions, but provides only weak confidence in the assignment of single individuals to coastal vs. elsewhere based on the genomic dataset.

A haplotype network of the mtDNA data (Figure 4 and Supporting Table 8) illustrates that only two haplotypes were present in HG—a result consistent with the findings of Sonsthagen et al. (2012). One of these was unique to HG and found in 5 out of 7 HG samples, and the other was found in a few individuals in Alaska (both AA and AK). The two haplotypes in HG were separated by a single mutational step, whereas the 19 haplotypes found elsewhere in North America were separated by up to four mutational steps.

**FIGURE 4.**
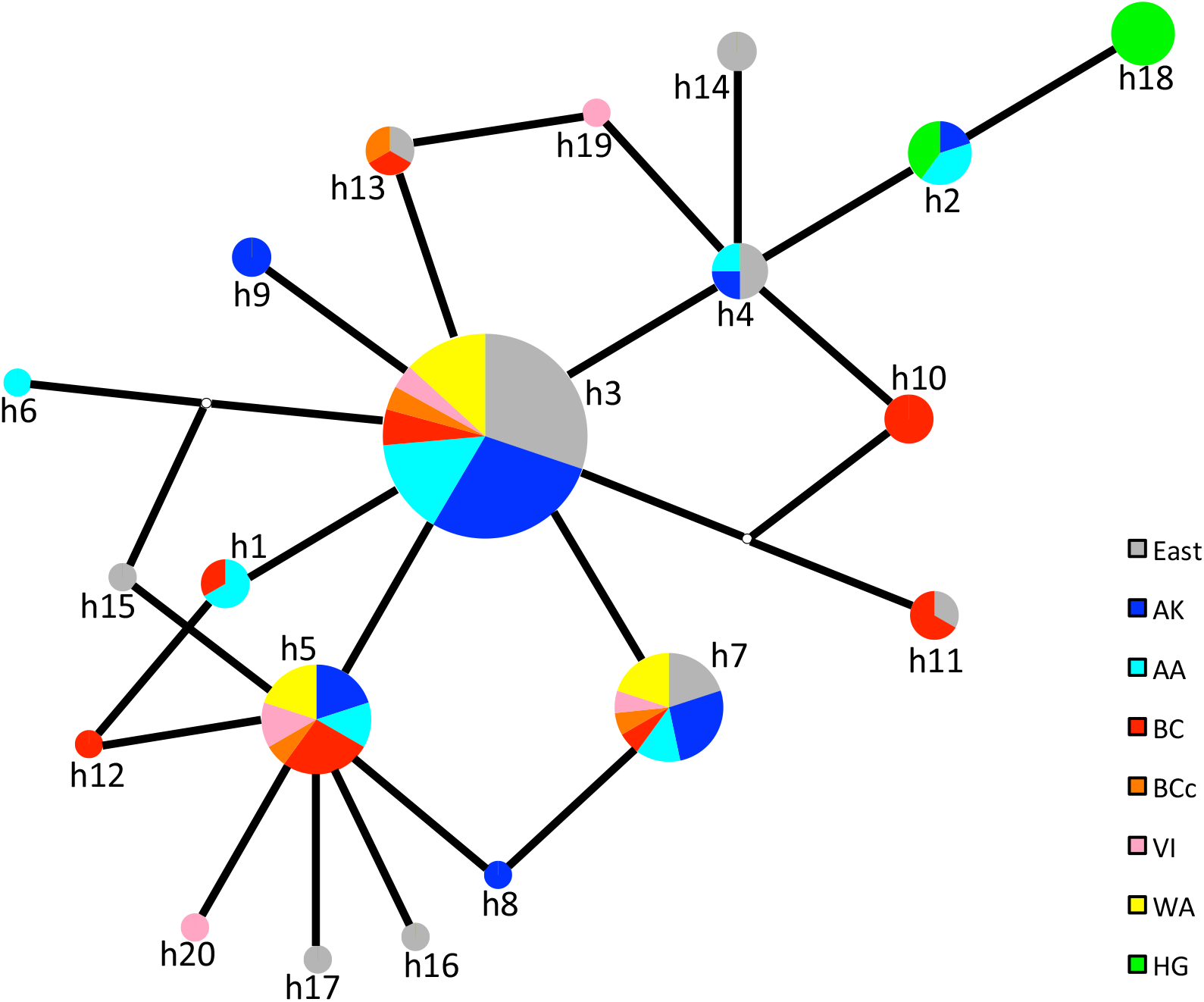
Median Joining network for mtDNA sequences (length 423 bp) obtained from 121 northern goshawks sampled in North America. Each pie chart represents a haplotype, with area proportional to its frequency in the sample set. Each black line indicates a single mutational step and small white dots indicate missing haplotypes. Population abbreviations can be found in Table 1, sample details in Supporting Table 1 and haplotype details in Supporting Table 8.

Estimates of genetic differentiation between sampling regions, both for the nuclear GBS and the mtDNA datasets, again reveal a pattern of clear differentiation of HG and little differentiation between other North American regions (Table 2). *F*_ST_ between HG and any of the other North American sampling regions ranges from 0.06 to 0.09 for the nuclear data and from 0.68 to 0.80 for the mtDNA data. Between any other North American sampling regions *F*_ST_ estimates are much lower, ranging from 0 to 0.01 for the nuclear data and from −0.12 and 0.08 for mtDNA. Inspection of the distribution of *F*_ST_ estimates for individual SNPs between HG and other North American goshawk samples (*n* = 9,850 SNPs in the North American dataset with MAF of 0.05 or higher) reveals that despite modest overall differentiation (weighted *F*_ST_ estimate is 0.115), there is considerable variation among SNPs with the top 1% of *F*_ST_ estimates being 0.736 or higher (Figure 5A and Supporting Table 5). In contrast, when HG is excluded the distribution of *F*_ST_ between areas currently considered within the range for *laingi* (i.e., VI, AA, and BCc) and areas considered within the range for *atricapillus* (i.e., BC, AK, WA, and East) is clustered much more tightly around zero (weighted *F*_ST_ estimate is 0.0059), with the threshold for the top 1% of *F*_ST_ estimates occurring at only 0.131 (Fig. 5B; *n* = 10,217 SNPs with MAF of 0.05 or higher).

**TABLE 2.** F_ST_ estimates between pairs of North American northern goshawk populations, based on the nuclear GBS dataset (weighted FST estimate of Weir & Cockerham, 1984) above the diagonal and the mtDNA dataset (sequence-based *F*_ST_ estimate of Hudson et al., 1992) below the diagonal. Population abbreviations and sample sizes are given in Table 1.

|  | HG | AA | AK | BC | BCc | East | VI | WA |
| --- | --- | --- | --- | --- | --- | --- | --- | --- |
| HG |  | 0.060 | 0.074 | 0.082 | 0.093 | 0.070 | 0.079 | 0.078 |
| AA | 0.708 |  | 0.006 | 0.007 | 0.010 | 0.006 | 0.013 | 0.005 |
| AK | 0.766 | -0.016 |  | 0.004 | 0.013 | 0.003 | 0.014 | 0.004 |
| BC | 0.676 | 0.061 | 0.075 |  | 0.007 | 0.002 | 0.012 | 0.004 |
| BCc | 0.747 | -0.044 | -0.083 | 0.017 |  | 0.000 | 0.011 | 0.007 |
| East | 0.724 | -0.022 | -0.006 | 0.042 | -0.057 |  | 0.007 | 0.002 |
| VI | 0.681 | 0.005 | -0.005 | -0.008 | -0.109 | 0.003 |  | 0.013 |
| WA | 0.804 | 0.014 | -0.031 | 0.077 | -0.123 | 0.012 | -0.037 |  |

**FIGURE 5.**
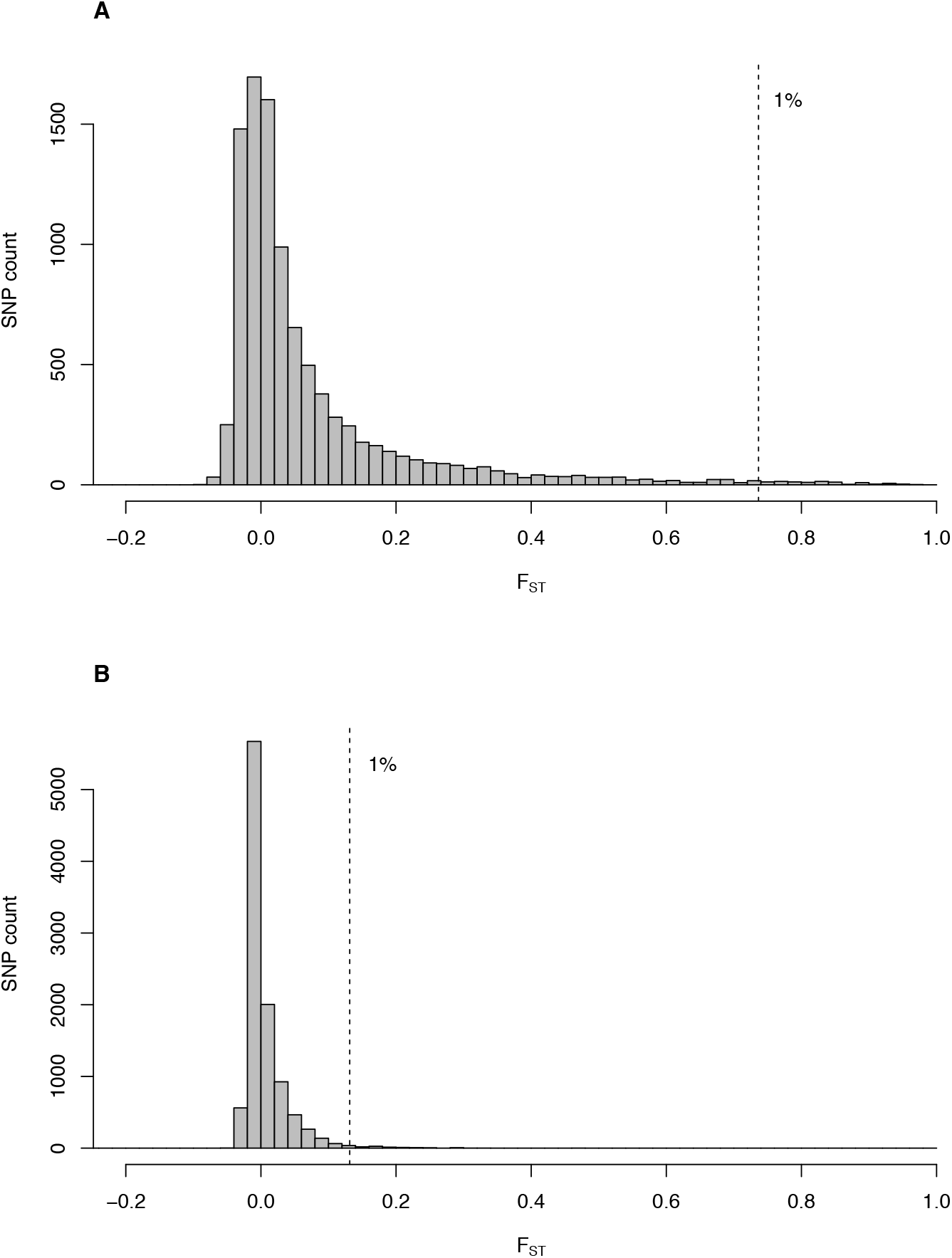
(A) Distribution among SNPs of genetic differentiation estimates between goshawks in Haida Gwaii and other areas of North America. Only SNPs with minor allele frequency of 0.05 or higher (*n* = 9,850) were considered. The dotted vertical line indicates where the top 1% of the *F*_ST_ distribution begins. (B) Similar to A, but excluding Haida Gwaii and comparing between areas often considered to be inhabited by *laingi* (i.e., the Alexander Archipelago, Vancouver Island, and the BC coast) and areas considered to be inhabited by *atricapillus* (i.e., interior BC, Washington state, Alaska north, and the eastern USA) (*n* = 10,217 SNPs).

### Genotyping of 10 informative loci

The addition of Taqman genotyping at ten loci allowed us to dramatically increase the number of individuals included in this study. With this extra information, we were able to more accurately map the geographic distributions of the two genetic clusters identified in the GBS study. To determine how closely this set of 10 loci recapitulates the GBS data (Figure 3), we compared the ancestry proportions calculated from these 10 loci with those estimated from the GBS dataset using the program Admixture. There is broad agreement between the two methods (Supporting Figure 9), except the admixture estimates tend to be somewhat higher when estimated using the subset of 10 loci. This discrepancy is likely due to lower precision of the 10-loci dataset compared to the thousands-of-loci GBS dataset.

For each sample we estimated an ancestry index (*S*) in HIest (Figure 6). All HG samples have ancestry indices above 0.5 (average *S* in HG is 0.92), whereas samples from elsewhere in North America have an average ancestry index of only 0.04 (with only one having an ancestry index of above 0.5). These results largely confirm, with a much larger sample size, the GBS finding that the HG genetic cluster is mainly confined to Haida Gwaii. Estimates of interclass heterozygosity can be informative with regards to the categories of admixed individuals. Interclass heterozygosity is expected to be very high in early hybrid generations and low in advanced hybrid generations. In our dataset, only four samples out of 386 (i.e., 1%) have interclass heterozygosity of 0.5 or higher suggestive of being first- or second-generation hybrids: two are from HG, one from AA and one from BCc (Supporting Figure 10).

**FIGURE 6.**
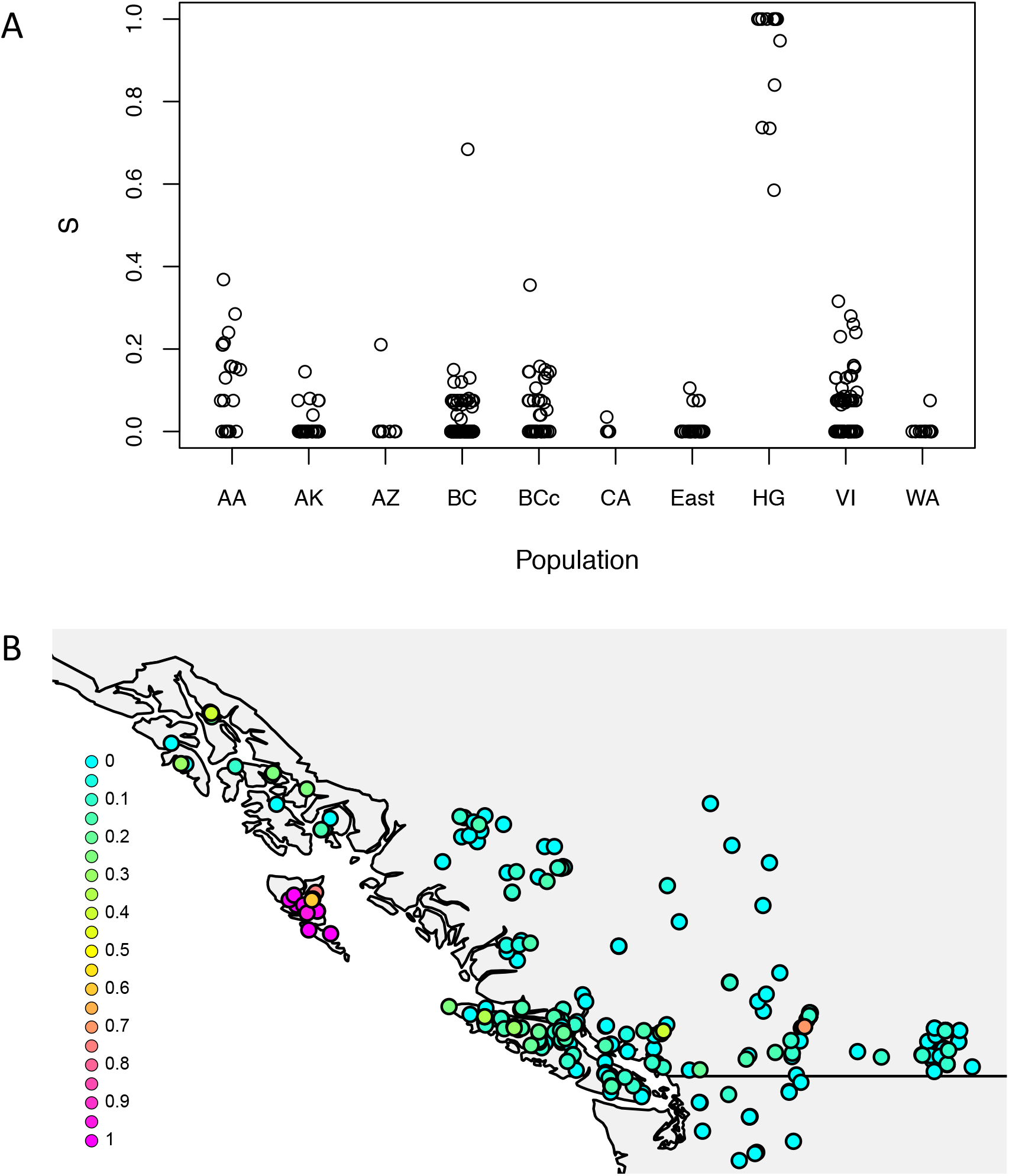
Estimates of ancestry proportions (*S*) from HIest based on 10 ancestry informative SNPs of northern goshawks for all North American Populations (A), with detail in (B) for the Pacific Northwest. Population codes are in Figure 1 and Table 1 and sample sizes in Table 1. Full results are in Supporting Table 6.

### Gene flow

While our results clearly indicate that HG is genetically distinct from other regions (Figures 3-6), there is some suggestion of recent gene flow between HG and other sampling regions, especially those coastal regions close to HG. Gene flow is suggested by the slightly closer similarity of these coastal regions to HG (i.e., AA, BCc, and VI) in the PCA (Fig. 3A) and the Admixture analysis (Fig. 3B), by the somewhat higher ancestry proportions (*S*) of some individuals from these regions (Fig. 6), as well as by the individuals that have high interclass heterozygosity (see above; Supporting Figure 10). Individuals with high interclass heterozygosity (*H* ≥ 0.5) are only found in HG, BCc, and AA; and individuals with intermediate interclass heterozygosity (0.25 ≥ *H* > 0.5) are additionally found in AK, BC interior, and VI.

To formally test for genetic mixture between genetically differentiated sampling regions we used Treemix with our GBS dataset to estimate the statistic f3. Four tests indicate admixture (significantly negative f3 test statistic), and in all four cases AA appears as a population that is admixed between HG and another sampling region: between HG and East (*Z* = −6.34520; *p* < 0.001), HG and WA (*Z* = −5.32896; *p* < 0.001), HG and BC (*Z* = −4.71371; *p* < 0.001), and HG and AK (*Z* = −4.69555; *p* < 0.001). The test statistic also suggests a trend for AA being admixed between HG and VI (*Z* = −1.41276; *p* = 0.079). Note that the f3 test only has power to reveal admixture when populations are differentiated.

### Timing of differentiation

To estimate the timing of the divergence between HG and remaining populations we used mutation rate estimates (see Methods) together with observed net nucleotide divergence (*D*_A_) between HG and other North American populations for both the nuclear and mtDNA datasets (Supporting Table 9). We advise caution regarding the resulting estimated divergence times because of the difficulty in applying long-term divergence rates to shorter time scales (due to saturation, for example) and because mutation rates may differ between *Ficedula* flycatchers and *Accipiter* hawks. Nonetheless, this approach can provide a very rough estimate of the timescale of the initial population separation between the Haida Gwaii and widespread North American clusters.

Resulting estimates of divergence between these genetic clusters are ~13 KYA for the mtDNA and ~24 KYA for the nuclear dataset. These results are about one order of magnitude lower than the estimated divergence time between North American and European goshawks, which is ~247 KYA for the mtDNA and ~346 KYA for the nuclear dataset.

## Discussion

Our analyses of variation in the nuclear and mitochondrial genomes of northern goshawks very clearly reveal a genetically distinct population in Haida Gwaii, in contrast to relatively subtle differentiation among other North American regions included in the study. Both principal component analysis and Admixture analysis of variation in over 6,000 nuclear SNPs from our set of high-quality DNA samples show clear genomic differentiation between northern goshawks from Haida Gwaii and those from all other North American sampling regions, including other coastal regions of British Columbia and Alaska. Only two mitochondrial haplotypes were found on Haida Gwaii, one of which was found just in the Haida Gwaii population. These two haplotypes are highly related compared to the haplotype variation seen within the rest of North America. Outside of Haida Gwaii, both nuclear and mitochondrial DNA show comparatively little geographic structure. However, there is some subtle differentiation in nuclear DNA signatures, with Pacific Northwest coastal regions (i.e., the Alexander Archipelago, the BC coast, and Vancouver Island) being somewhat differentiated from other parts of North America.

We clarified the ranges of the two main genetic clusters by genotyping a larger set of individuals (386) using 10 informative loci that have large frequency differences between the clusters. This approach confirmed that the range of one genetic cluster is almost entirely restricted to Haida Gwaii, whereas the other genetic cluster encompasses almost all individuals from other sampling regions in North America, including those along the west coast such as Vancouver Island and southeast Alaska. One notable exception to this finding is a single sample from Vernon, B.C., approximately 950 km from Haida Gwaii, which has a high fraction of alleles common in Haida Gwaii. This specimen may represent a long-distance dispersal and admixture event.

We are not the first to document genetically distinct forms of taxa on Haida Gwaii. Pruett et al. (2013) summarized genetic evidence for a Haida Gwaii glacial refugium in diverse taxa, from plants to mammals, fishes, and birds. Among the 11 bird species they surveyed, fully seven exhibited genetic signals of long-term occupancy of Haida Gwaii, including four with endemic subspecies. The life histories of these refugial populations suggest the presence of a forested refugium, and our results in the northern goshawk add support to that inference.

Our genetic results are not concordant with the prevailing understanding of the distributions of the two subspecies *A. g. laingi* and *A. g. atricapillus* in British Columbia and southeast Alaska. The prevailing taxonomic treatment of goshawks in this region might suggest two genetic clusters, perhaps with some intergradation between them, corresponding closely to the subspecies ranges as defined through morphological analysis. The presently accepted understanding of the distribution of the subspecies *A. g. laingi* (COSEWIC 2013) is based on Taverner’s (1940) description of that dark-coloured form occurring in Haida Gwaii, along with evidence that somewhat dark individuals can also be found on Vancouver Island (Taverner, 1940) and the Alexander Archipelago of southeast Alaska (Webster 1988, Titus et al., 1994). These morphological observations were supplemented by habitat modelling to infer the currently mapped range of *A. g. laingi* as presented by COSEWIC (2013), which includes much of coastal BC, southeast Alaska, and the western part of Washington state. In contrast, our genomic analysis reveals two clearly differentiated genetic clusters, but these correspond to Haida Gwaii versus elsewhere. Within the non-Haida Gwaii cluster, there is some subtle differentiation between sites traditionally considered within the *laingi* range (Vancouver Island, southeast Alaska, and the BC coast) and elsewhere (i.e., within the traditional *atricapillus* range), but this differentiation is small compared to the very clear and diagnosable difference between Haida Gwaii and elsewhere. Interestingly, much of the variation among the non-Haida Gwaii populations is orthogonal to that between Haida Gwaii and the other populations (Fig. 3), suggesting differentiation among the other populations is not attributable solely to gene flow with Haida Gwaii. We note that the subtle patterns of differentiation do not appear to be completely explained by a simple isolation-by-distance model, since Alaska north and the eastern USA have very similar genomic signatures in contrast to the slight differentiation between those areas and southeast Alaska, Vancouver Island, and the BC coast. This pattern may in part be due to multiple range expansions following Pleistocene glaciations, and possibly also by patterns of local adaptation.

While the two differentiated genetic clusters of goshawks in British Columbia indicate some degree of restricted gene flow and independent evolution, the dark-plumage variant shared between goshawks on Haida Gwaii and at least some goshawks in other coastal regions of BC and southeast Alaska may be a shared adaptation to coastal conditions, possibly enhanced by some level of shared ancestry and gene flow among these regions. This general trait of darker plumage occurs among other avian subspecies in this area (e.g., *Ardea herodias fannini*, *Accipiter striatus perobscurus*, *Aegolius acadicus brooksi*, and *Picoides villosus picoideus*). In two of these cases, a named subspecies endemic to Haida Gwaii is both darker and genetically differentiated from populations elsewhere: Northern Saw-Whet Owl (*Aegolius acadicus brooksi*; Withrow et al., 2014) and Hairy Woodpecker (*Picoides villosus picoideus*; Topp & Winker 2008; Klicka et al., 2011). The simultaneous occurrence among multiple independent lineages suggests that darker colour has adaptive value, and is consistent with Gloger’s rule (Gloger 1833), a tendency for endotherms to be darker in humid environments such as the coastal climate of this region (e.g., Burtt & Ichida, 2004).

As noted, the dark plumage characteristic upon which the subspecies *laingi* is based has a wider apparent distribution than just Haida Gwaii, but a pattern of apparent phenotypic intergradation with the more widespread *atricapillus* outside of Haida Gwaii has been known since the original description (Taverner, 1940; Webster, 1988; Titus et al., 1994). Taverner’s (1940: p. 160) description of the *laingi* phenotype noted that it was “most typical on the Queen Charlotte Islands, the birds of Vancouver Island being more variable and less plainly characterized.” He did not examine material from southeast Alaska. Webster (1988: p. 46) and Titus et al. (1994) did, and both concluded that coloration of birds from southeast Alaska was quite variable, such that the area could be viewed as a mix of *laingi* and *atricapillus* phenotypes. Webster (1988) noted that the darkest southeastern Alaska specimens “are not quite as black as those from the Queen Charlotte Islands, but just as dark as those from Vancouver Island.” Our genetic data reflect this phenotypic intergradation to some degree (Fig. 6), with southeast Alaska and coastal regions of BC showing slightly more genetic similarity (compared to other North American regions) to Haida Gwaii. However, the genomic data contrast with the phenotypic patterns in showing that southeast Alaska and Vancouver Island are much more similar genomically to the widespread North American genetic cluster than they are to the Haida Gwaii genetic cluster.

Our estimated divergence time between the Haida Gwaii and widespread North American genetic clusters (~13 KYA for the mtDNA and ~24 KYA for the nuclear dataset) corresponds somewhat well with the end of the last glacial maximum (roughly 20,000 years ago; Yokoyama et al., 2000; Clark et al., 2009). While much of inland British Columbia was covered with glaciers at that time, there is much evidence that areas along the coast including Haida Gwaii and the Alexander Archipelago (in southeast Alaska) had large ice-free areas of refugial habitat that were used by a variety of species (Shafer et al., 2010). Moreover, Haida Gwaii was connected to the mainland due to the lower sea level, and the exposed Hecate Strait had areas of forest (Lacourse et al., 2003) that presumably could have been inhabited by goshawks. A plausible scenario is that northern goshawks were distributed across these coastal areas during glacial periods, and the subsequent melting of glaciers and rise of sea levels resulted in separation of the Haida Gwaii population from the mainland population. This scenario has been proposed for a variety of other bird populations on Haida Gwaii (Pruett et al., 2013).

Our data suggest that the Haida Gwaii population of goshawks has been moderately isolated from other parts of North America since that rise in sea levels. However, some population connectivity is suggested by the Treemix analysis, the PCA, and *F*_ST_ estimates, each of which suggests some occasional gene flow between Haida Gwaii and the Alexander Archipelago (and perhaps other nearby areas such as Vancouver Island or the BC coast). Furthermore, we did find two Haida Gwaii individuals in our genotype assay dataset that were supported as “*laingi* backcrosses”. This finding lends some support to the idea that there have been very recent dispersal events to Haida Gwaii (followed by interbreeding), but there is some uncertainty in this assessment due to the small number of loci genotyped in those individuals.

In managing any species of conservation concern, understanding its range and its genetic relationships with other populations is important to effective management. Currently listed as Threatened under the Canadian Species at Risk Act, the range of *laingi* has been considered to encompass Haida Gwaii, the Alexander Archipelago, Vancouver Island, the BC coast, and coastal Washington. Within this currently understood range the population of goshawks was estimated to be ~1200 individuals (COSEWIC, 2013), and conservation policy is currently tailored based on this population size and range. Our results suggest a taxonomic re-evaluation of the *laingi* subspecies may be appropriate, but this would likely depend in part on more detailed morphological analysis beyond the scope of the present paper. We note however that the subspecies concept is the subject of much debate, particularly in terms of how much subspecies taxonomy should depend on genetic clustering vs. specific morphological traits contained in the initial subspecies description (Winker, 2009, 2010; Patten, 2010; Liu et al., 2018). Regardless of taxonomic treatment, we think the evidence in support of treating the Haida Gwaii population as a distinct conservation unit (e.g., a “designatable unit”) is strong: Haida Gwaii goshawks are very clearly genetically distinct, have a recognized phenotype (darker plumage than elsewhere, even compared to those in southeast Alaska and Vancouver Island), and inhabit an ecologically differentiated and geographically separated archipelago. This Haida Gwaii population is presumably at very high risk of extinction, given its extremely small (and historically declining) size of just 48-57 individuals (COSEWIC, 2013).

While the Haida Gwaii population size is thought to have declined from historical levels (COSEWIC 2000, 2013), it was likely never very large, given the typical territory sizes of northern goshawks and the limited size of the archipelago. Small populations are of special conservation concern because they tend to have low diversity, to experience inbreeding, and to respond poorly to natural selection given the power of genetic drift on small populations. Our data provide some insights into these three typical characteristics of small populations. First, levels of nuclear nucleotide variability in Haida Gwaii are about 80% of that in other North American sampling regions (Supporting Table 10). This reasonably high variability (given the small geographic range and population size) is likely a combined result of the relatively recent (i.e. within the last 20,000 years or so) population differentiation between Haida Gwaii and elsewhere and the occasional gene flow from other regions to Haida Gwaii. Second, our data do not reveal a higher amount of inbreeding in Haida Gwaii, as inbreeding coefficients range from −0.0621 to 0.278 which is within the range in other populations (Supporting Table 11 and Supporting Figure 11) and consistent with previous estimates (Sonsthagen et al., 2012). This suggests inbreeding is not presently a major concern. However, further decline and/or isolation of the population might elevate the possibility of inbreeding depression. Third, while genetic drift certainly is expected to overpower weak selection in a small population, our highly skewed distribution of *F*_ST_ values in the comparison of Haida Gwaii and elsewhere suggests that strong selection has likely shaped some characteristics of the Haida Gwaii population. Future research will more specifically test for the role of selection in shaping patterns of genomic variation in these populations.

Outside of Haida Gwaii, populations of goshawks in British Columbia and nearby regions have also declined from historical levels. This decline has been particularly closely examined for regions currently considered to be inhabited by *laingi*, given the listing of that taxon as Threatened (COSEWIC 2000, 2013). It should also be noted the *atricapillus* subspecies (as currently delineated) was recently designated as Blue Listed by the British Columbian government due in large part to a precipitous ~95% population decline observed in interior central and northwestern populations (B.C. Conservation Data Centre 2017; Doyle et al., 2017). Blue List status in British Columbia indicates that this organism is of special conservation concern within the province. Our genetic results have two implications for management of goshawk populations outside of Haida Gwaii: First, there is only relatively subtle differentiation between coastal populations (e.g., Vancouver Island, southeast Alaska, and the BC coast) and BC Interior populations, suggesting that an integrated approach to management may be most effective. Second, the subtle differentiation that is observed between some sampling regions suggests that goshawks tend to have limited gene flow between regions; this suggests that the causes of observed regional declines (Doyle et al., 2017) may be somewhat specific to each region and management strategies tailored to each region may be necessary to maintain local populations.

While the overall decline of goshawk populations in British Columbia and surrounding regions deserves attention, the Haida Gwaii population merits particularly urgent focus. With its small population size, the genomically distinct Haida Gwaii population can be considered to be one of the most endangered organisms on the planet. The Haida Gwaii population represents the core range of the legally Threatened *laingi* subspecies (as currently defined) of northern goshawk and is reasonably isolated from other goshawk populations, which are differentiated genomically. The current population size estimate of just 48-57 mature individuals (COSEWIC, 2013) means that, if it is considered a Designatable Unit under the Canadian Species at Risk Act, it would meet Endangered status based on population size alone (< 250 mature individuals). Given the continued threat of habitat loss and conflict with humans, this population is clearly at extreme risk for extinction and conservation efforts for this population should reflect the significance of losing a unique large vertebrate predator forever.

## Acknowledgements

For providing funding to this research we thank Genome British Columbia, British Columbia Ministry of Forests, Lands and Natural Resource Operations, Coast Forest Products Association, and Western Forest Products Inc. (all partners in the Genome BC User Partnership Program Grant UPP023), and the Natural Sciences and Engineering Research Council of Canada (Discovery grant RGPIN-2017-03919 to DEI). For advice during project planning we especially thank Steve Gordon, John Deal, and Bryce Bancroft. For providing valuable support through sample contribution and/or advice regarding this project we thank A&A Trading Ltd., the NCE Sustainable Forest Management Network, Forest Science Program of the BC Forest Investment Account, Island Timberlands, Tembec Inc, the School of Natural Resources and the Environment at The University of Arizona, Sally Aitken, Janice Anderson, Bryce Bancroft, Carita Bergman, Sharon Birks, Victoria Bowes, Melanie Bucci, Christina Carrières, Melanie Culver, John Deal, Kiku Dhanwant, Steve Gordon, Janet Hinshaw, Molly Hudson, Jessica Irwin, Jeremy Kirchman, Karl Larsen, Todd Mahon, Ben Marks, Brent Matsuda, Sue McDonald, Erica McClaren, Kathy Molina, Gerry Morigeau, Jacques Morin, Loren Rieseberg, Dolph Schluter, Kari Stuart-Smith, Ildiko Szabo, Sandy Talbot, Tania Tripp, Thomas Trombone, Thijs Valkenburg, Carla Vargas, Martina Versteeg, Warren Warttig, Berry Wijdeven, Melanie Wilson, the American Museum of Natural History, Beaty Biodiversity Museum, Burke Museum of Natural History and Culture, Field Museum of Natural History Bird Collection, New York State Museum, O.W.L. (Orphaned Wildlife) Rehabilitation Society, UCLA Dickey Collection of Birds and Mammals, the University of Alaska Museum, the University of Arizona Natural History Museum, and the University of Michigan Zoological Collections.

